# Freely chosen cadence during cycling attenuates intracortical inhibition and increases intracortical facilitation compared to a similar fixed cadence

**DOI:** 10.1101/2020.04.23.056820

**Authors:** Simranjit Sidhu, Benedikt Lauber

**Affiliations:** Adelaide Medical School, The University of Adelaide, Adelaide, Australia; Department of Neuroscience and Movement Science, University of Fribourg, Fribourg, Switzerland; Department of Sport and Sport Science, University of Freiburg, Freiburg, Germany

**Keywords:** cycling cadence, intracortical inhibition, intracortical facilitation, motor cortex

## Abstract

In contrast to other rhythmic tasks such as running, the preferred movement rate in cycling does not minimize energy consumption. It is possible that neurophysiological mechanisms contribute to the choice of cadence, however this phenomenon is not well understood. Eleven participants cycled at a fixed workload of 125 W and different cadences including a freely chosen cadence (FCC, ∼72), and fixed cadences of 70, 80, 90 and 100 revolutions per minute (rpm) during which transcranial magnetic stimulation (TMS) was used to measure short interval intracortical inhibition (SICI) and intracortical facilitation (ICF). There was significant increase in SICI at 70 (*P* = 0.004), 80 (*P* = 0.008) and 100 rpm (*P* = 0.041) compared to FCC. ICF was significantly reduced at 70 rpm compared to FCC (*P* = 0.04). Inhibition-excitation ratio (SICI divided by ICF) declined (*P* = 0.014) with an increase in cadence. The results demonstrate that SICI is attenuated during FCC compared to fixed cadences. The outcomes suggest that the attenuation of intracortical inhibition and augmentation of ICF may be a contributing factor for FCC.

## Introduction

Compared to other rhythmic motor tasks such as walking and running, the preferred cycling cadence at a constant workload does not minimize energy consumption (i.e. freely chosen cadence; FCC) (Marsh & Martin, 1993). The factors underlying this effect are not well understood, but neural mechanisms might contribute. There is some evidence to suggest that rhythmical locomotor movements such as cycling are mediated by not only spinal (Capaday *et al*., 1999; Petersen *et al*., 2003; Zehr *et al*., 2009) but also supraspinal mechanisms (Petersen *et al*., 2001a; Sidhu *et al*., 2012; Forman *et al*., 2014). Using positron emission tomography (PET), bilateral increase in activation of the primary motor cortex (M1) and primary somatosensory cortex during active cycling has been reported (Christensen *et al*., 2000). Furthermore, the activation of M1 was positively correlated with an increase in cadence, suggesting increased levels of brain activation with an increase in cycling cadence. Similarly, with the use of functional magnetic resonance imagining (fMRI), an increase in activation of both motor cortices with an increase in pedaling frequency has been demonstrated (Mehta *et al*., 2012); further supporting the notion that motor cortical activity is augmented with increased pedaling frequency. While imaging has been fundamental in identifying the neural networks and regions of interest during locomotion, a more refined understanding of the underlying neural mechanisms using non-invasive brain stimulation techniques is warranted to determine whether these brain areas are directly contributing to locomotion. This information can be easily obtained by directly asses the activity in the cortical areas concerned, such as by the use of transcranial magnetic stimulation (TMS).

A recent study investigated the effects of cadence on spinal and supraspinal neural circuity during arm cycling by applying TMS on the motor cortex and by electrically stimulating the corticospinal tract at 30, 60 and 90 repetitions per minute (rpm, Forman *et al*., 2015). They reported enhanced corticospinal excitability, both at cortical and spinal levels during the elbow flexion phase. Interestingly, during the elbow extension phase, corticospinal excitability was increased but spinal excitability was reduced. It however remains unclear as to whether the increase in corticospinal excitability during the extension phase (Forman *et al*., 2015) and the increase in levels of brain activity reported with imaging (Mehta *et al*., 2012) is due to a modifications in excitability of GABAergic inhibitory circuits in layer 1-3 of the motor cortex (Di Lazzaro & Rothwell, 2014) and/or modulations in the excitability of NMDA receptor networks (Ziemann, 2013). One way to test for inhibition and/or excitation within the motor cortex is via the application of paired-pulse TMS over the area representing the muscle of interest, while measuring the evoked responses recorded via surface electromyography (i.e. motor evoked potential; MEP). In paired-pulse stimulation, a conditioning TMS pulse precedes a test pulse by several milliseconds. Depending on the interstimulus interval (ISI) between the two TMS pulses, the test pulse response becomes either smaller – testing for short interval intracortical inhibition (SICI, ISIs between 2–5 ms) (Lazzaro *et al*., 1998; Di Lazzaro *et al*., 2001) or larger– testing for intracortical facilitation (ICF, ISIs between 7–20 ms) relative to an unconditioned single pulse response (Ziemann *et al*., 1996). While most previous studies have used arm cycling as a model to study the neural control of cycling, the present study investigated the influence of cadence on motor cortical inhibition and facilitation during leg cycling. More specifically, the influence of FCC versus a comparable prescribed cadence on intracortical circuitry remains undetermined. The primary aim of the present study was to investigate the influence of FCC on the excitability of the inhibitory versus excitatory networks in M1. Given that humans tend to pick a cadence during leg cycling that is not energetically optimal, it is theorised that brain mechanisms, and more specifically increased cortical output, may contribute to this effect. Therefore, it was hypothesized that SICI would be attenuated at FCC while ICF would be augmented at FCC compared to a similar fixed cadence.

## Methods

### Participants

Eleven young healthy active subjects (25.9 ± 3.8 years, 8 males; 76.4 ± 15.7 kg; 176.8 ± 8.7 cm) participated in the study. All subjects were sport and exercise science students and recreationally active. Participants gave written informed consent before the start of the experiment and did not present with any neurological or cardiovascular disorders. The experiment was approved by the Ethics committee of the University of Freiburg and was in accordance with the declaration of Helsinki.

### Experimental procedures

After giving informed consent and subject preparation, the optimal position on the motor cortex for TMS application was established, while subjects were seated on the bike. Subjects were then asked to cycle at a freely chosen cadence at 125 watts (W) for 5 minutes to warm up and familiarise themselves to the cycling workload. Subsequently, subject cycled another 5 minutes without cadence feedback to establish their FCC. This was followed by a further 3 min cycling where the AMT was established. To determine the FCC and cadence dependent effect on intracortical mechanisms, subjects cycled at the various cadences while TMS was applied.

### Cycling

Subjects cycled on a cycle ergometer (Ergobike medical 8, Daum electronic GmbH, Fuerth, Germany) at a constant workload of 125 W. Subjects started with FCC since pilot testing revealed that the FCC varied depending on the cadence subjects cycled at in a previous trial; whereby subjects tended to cycle at higher cadences when the preceding cadence was high and at lower cadences when the preceding cadence was low. The FCC was established while subjects cycled for 5 minutes without feedback of their cadence. An average of the cadence from minutes 2-5 was taken as the FCC. Subjects were then instructed to cycle at their FCC (group mean FCC: 71.6 ± 8.1 rpm) with the cadence visually displayed in front of them. Following which, the other prescribed cadences including 70, 80, 90 and 100 rpm were tested in a randomized order. At the beginning of each trial, subjects were instructed to cycle at the defined cadence for 2 minutes with visual feedback of their cadence before any stimulations were given.

### Electromyography (EMG)

Surface electromyographic (EMG) recordings were taken from the vastus lateralis (VL) of the left leg. The skin was shaved and cleaned with alcohol swabs before surface EMG electrodes (Blue sensor P, Ambu, Bad Nauheim, Germany) were attached according to SENIAM guidelines with an inter-electrode distance of 2 cm. The EMG recordings were amplified (x 500), bandpass filtered (10–1000 Hz), and sampled at 4kHz. Subjects cycled for approximately 3 minutes at 125 W while the VL EMG was recorded. All data was stored on a computer using a custom-built software (LabView based, National Instruments, Austin, TX) for off-line analysis. Maximal cycling EMG in the VL was calculated as the maximal rectified EMG activity observed at each cadence. This was also done to determine the position on the crank cycle for delivery of the stimulations which coincided with 50% of the maximum rectified cycling EMG (Sidhu *et al*., 2013). As the aim of the study was to only look at cadence effects during extension phase of cycling, we only recorded EMG from the VL and not from antagonistic muscles.

### Transcranial Magnetic Stimulation (TMS)

TMS was applied on the right primary motor cortex using a Magstim® 200^2^ stimulator with a Bistim unit using a double cone coil (Magstim®, Whitland, UK) to activate the left quadriceps. Brainsight TMS navigation (Brainsight 2®, Rogue Research, Montreal, Canada) was used to track the position of the coil relative to the skull ensuring that the defined coil position remained constant throughout all stimulations. The optimal site for evoking MEPs in the VL was determined by initially setting the starting point at 0.5 cm anterior to the vertex, and over the midline. The final coil position was established by moving the coil anterior and right from the vertex resulting in the greatest MEP inducing a posterior-anterior flow of current. During cycling, TMS was triggered at 50% of the rising phase of the rectified maximal EMG (Sidhu *et al*., 2013; Figure 1). The stimulation position was established during the freely chosen cadence trial and was kept constant at all other cadences. The level of muscle activity was similar between cadences at time of stimulation.

**Figure 1.**
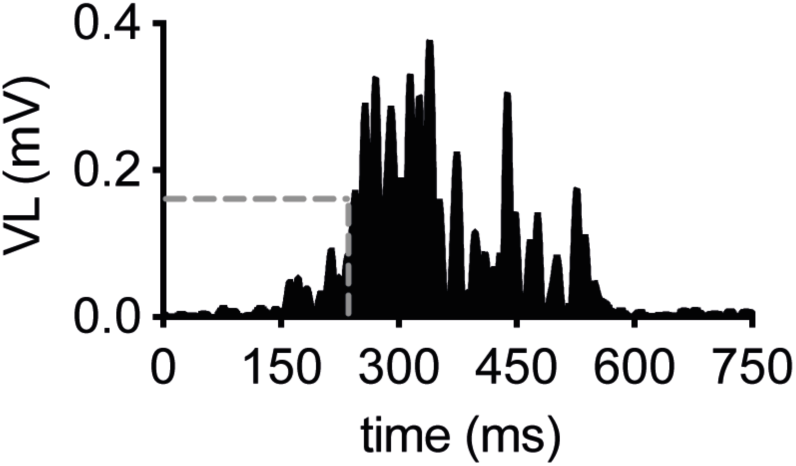
Averaged EMG from a subject. Dashed lines show the selected threshold for the TMS stimulation. This threshold was at 50% of the maximum rectified cycling EMG during the activation phases of the VL muscle.

### Active Motor Threshold (AMT)

Active motor threshold was defined as a clearly visible MEP response larger than the background EMG while subjects were cycling at a constant workload of 125W during FCC. Active motor threshold was 39 ± 7 % of maximal stimulator output.

### Short-interval intracortical inhibition

Short interval intracortical inhibition (SICI) was measured by applying two TMS pulses with an interstimulus interval of 2.5 ms (Wälchli *et al*., 2017; Lauber *et al*., 2018) where the first conditioning pulse was subthreshold (0.7 AMT) and the second test pulse was suprathreshold (1.2 AMT). The conditioning pulse activates intracortical inhibitory interneurons and attenuates the MEP evoked by the second pulse which reflect excitability of the corticospinal neurons (Di Lazzaro & Rothwell, 2014).

### Intracortical Facilitation

Intracortical facilitation (ICF) was measured in the same way as SICI, except that the ISI between the conditioning and the test pulse was 10 ms since consecutive stimuli applied at this interval causes a facilitation of the test pulse response (Soto *et al*., 2006).

### Stimulation Protocol

A total of 60 stimulations (20 single pulses, 20 paired pulses to measure SICI and 20 paired pulses to measure ICF) were applied at each cadence with a random interval of between 4-6 seconds. As expected, the crank position at which the TMS was given varied slightly, but not significantly between cadences: 70 rpm: 265.2 ± 7.9 deg; 80 rpm: 262.2 ± 9.9 deg; 90 rpm: 260.8 ± 6.3 deg; 100 rpm: 254.5 ± 9.4 deg; FCC: 272.8 ± 9.7 deg (from top dead center). Crank position was monitored by a light barrier which was mounted to the frame of the ergometer. The light barrier counted the number of teeth (52) of the chain ring during each crank cycle such that the crank angle could be calculated with an accuracy of 6.9 degrees.

### Data analyses and Statistics

All data was analysed using custom written Matlab scripts (Mathworks Inc., Chatswool, MA, USA).

### Motor Cortical Inhibition (SICI) and Facilitation (ICF)

The size of the MEP was quantified by measuring the peak to peak amplitude. The peak to peak amplitude of the conditioned MEP was compared to an unconditioned MEP. SICI was expressed as percentage inhibition of the conditioned MEP compared to an unconditioned MEP using the formula: 100 – (conditioned MEP/ unconditioned MEP × 100). ICF was quantified as the percentage facilitation of the conditioned MEP in relation to the unconditioned MEP according to the formula: (conditioned MEP/test MEP × 100) – 100 (Kuhn *et al*., 2016; Lauber *et al*., 2018).

### Inhibition-Excitation Ratio

In order to evaluate the relationship between inhibition (SICI) and excitation (ICF), inhibition-excitation ratios (I/E ratio) were calculated by dividing the average amount of SICI with the average amount of ICF.

### Corticospinal Excitability

Corticospinal excitability was quantified as the peak to peak amplitude of the unconditioned MEP elicited with single pulse TMS.

### Maximal cycling EMG activity

Muscle activity during cycling was evaluated by calculating root mean square values of the EMG of 20 full crank cycles (on and off phase) without TMS. The maximal EMG value in each of these cycles was then measured and averaged across the 20 trials.

### Background EMG Activity (bEMG)

Muscle activation at the time of the TMS was calculated as root mean square over a short timeframe prior to the TMS. The length of this timeframe was adjusted individually for each cadence representing 5% of the total time taken for one crank revolution (Forman *et al*., 2015). Accordingly, the windows were 70 rpm: 42.85 ms, 80 rpm: 37.5 ms, 90 rpm: 33.3ms, 100 rpm: 30 ms, FCC: 44.18 ± 5.3 ms.

### Cycling Performance

Cycling performance was based on how constant the subjects cycled at each cadence and by calculating the coefficient of variation of the pedaling rate.

### Statistics

Normal distribution of the data was checked using the Shapiro-Wilk test. As normal distribution was violated even after log transformation, non-parametric Friedman ANOVA was used to identify difference in SICI, ICF, I/E ratios, MEPs and background EMG across cadences. When ANOVAs revealed significant main effect of cadence, Bonferroni-corrected post hoc tests were used to identify the difference between the cadences. Effect sizes were calculated for significant results by Kendall’s W (r). Differences between the conditioned and unconditioned MEP (for SICI and ICF) were tested using Wilcoxon signed-rank tests and Bonferroni corrected t-tests. Because cycling EMG, cycling performance and crank position were normally distributed, one-way ANOVA followed by Bonferroni t-tests corrected for multiple comparisons were used to calculate differences between cadences. As TMS measurements, especially under dynamic conditions can be variable, coefficient of variation was calculated for all TMS pulses. All data are reported as means ± standard deviation (SD). SPSS 24 (Chicago, IL, USA) software was used for all statistical comparisons and α level was set at 5% (*P* ≤ 0.05).

## Results

### SICI

At all cadences, there was significant inhibition of the conditioned MEP compared to the unconditioned MEP (Table 1). There was a significant effect of cadence on SICI (χ^2.^ = 10.40, df = 4, *P* = 0.034). SICI was greater at 70 (*P = 0*.*004, r = 0*.*47*), 80 (*P = 0*.*008, r = 0*.*60*) and 100 rpm (*P = 0*.*041, r = 0*.*49*) compared to FCC but not at 90 rpm (*P* = 0.07, r = 0.36, Fig. 2, 4A).

**Table 1.** shows the SICI and ICF results at each of the cadences along with the corresponding statistical results.

| Cadence | SICI (mean ± SD) | *Wilcoxon-Test* | ICF (mean ± SD) | *Wilcoxon-Test* |
| --- | --- | --- | --- | --- |
| 70rpm | 23.2 ± 4.1 | $p = 0.017$ | 10.6 ± 2.7 | $p = 0.007$ |
| 80rpm | 22.6 ± 3.9 | $p = 0.0007$ | 15.7 ± 3.7 | $p = 0.013$ |
| 90rpm | 19.6 ± 4.3 | $p = 0.010$ | 20.1 ± 3.1 | $p = 0.004$ |
| 100rpm | 20.6 ± 2.9 | $p = 0.0002$ | 24.9 ± 3.9 | $p = 0.019$ |
| FCC | 11.5 ± 2.3 | $p = 0.002$ | 21.9 ± 3.5 | $p = 0.048$ |

**Figure 2.**
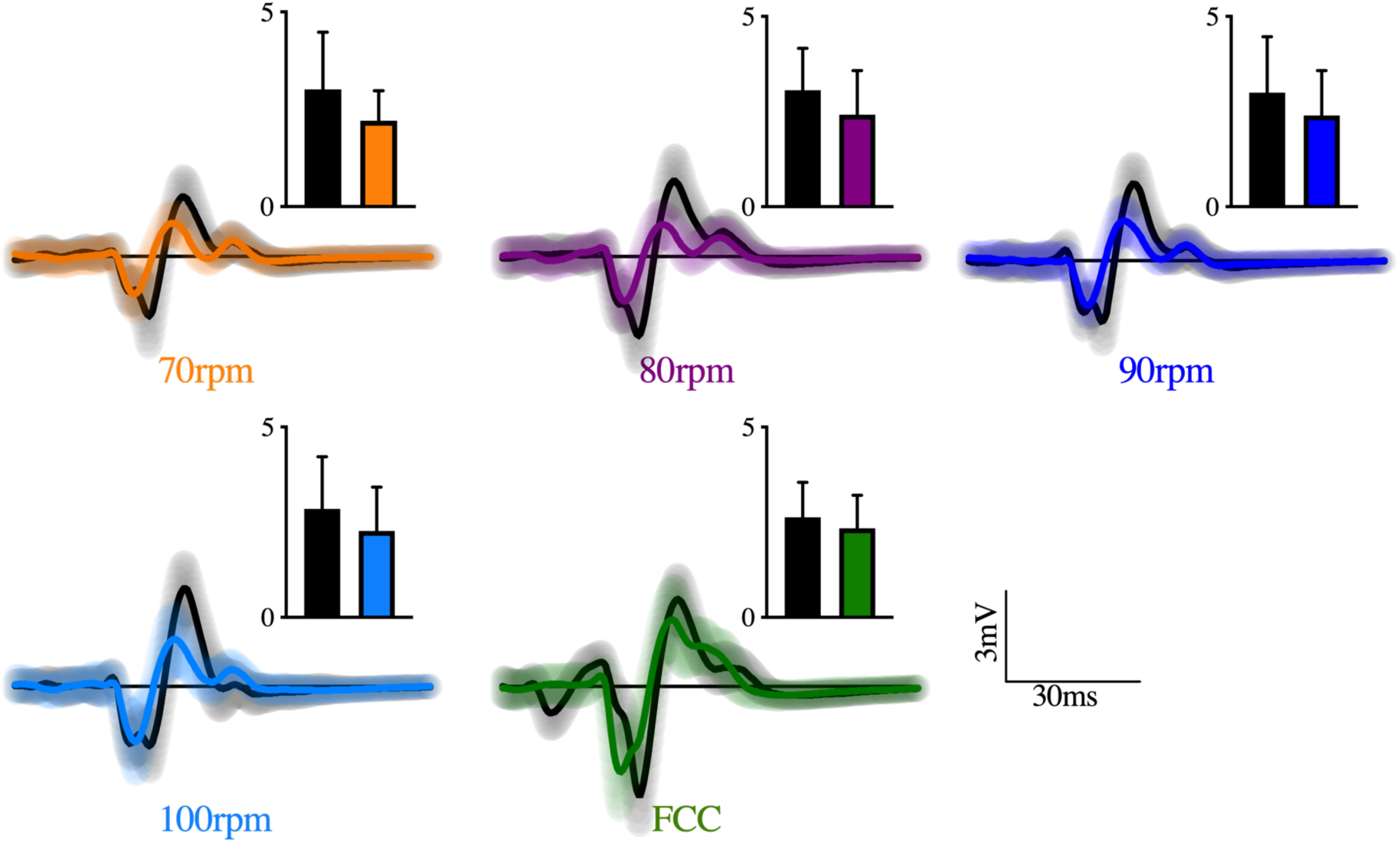
Representative TMS responses (SICI) from a subject at each of the cadences. The black lines represent the unconditioned MEPs while the colored lines show the results from the conditioned MEP (SICI). The black bars show the mean peak to peak amplitudes (+SD) of the conditioned MEP (SICI, colored bar) and the unconditioned MEP (black bar).

**Figure 3.**
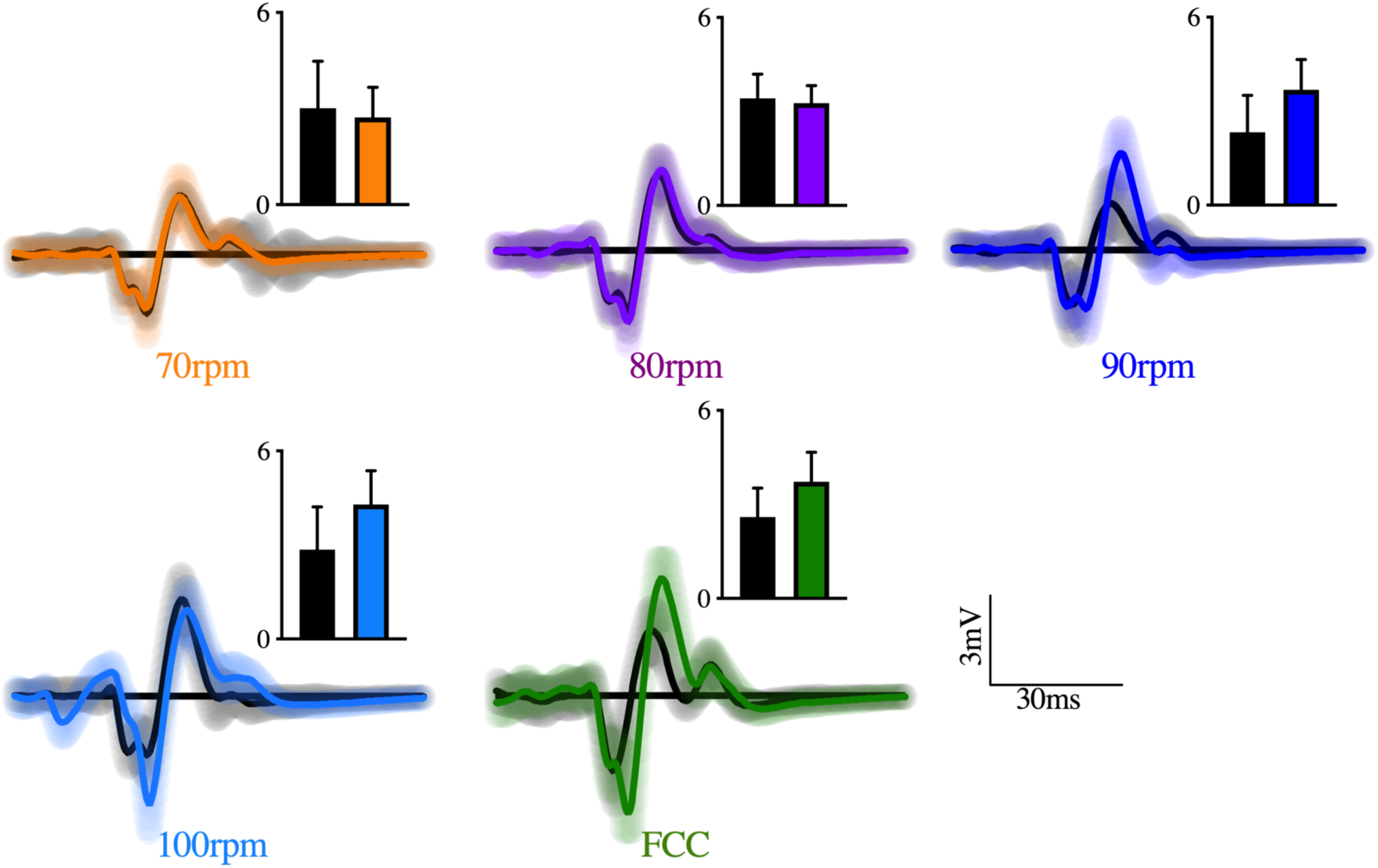
Representative TMS responses (ICF) from a subject at each of the cadences. The black lines represent the unconditioned MEPs while the coloured lines show the results from the conditioned MEP (ICF). The black bars show the mean peak to peak amplitudes (+SD) of the conditioned MEP (ICF, colored bar) and the unconditioned MEP (black bar).

**Figure 4.**
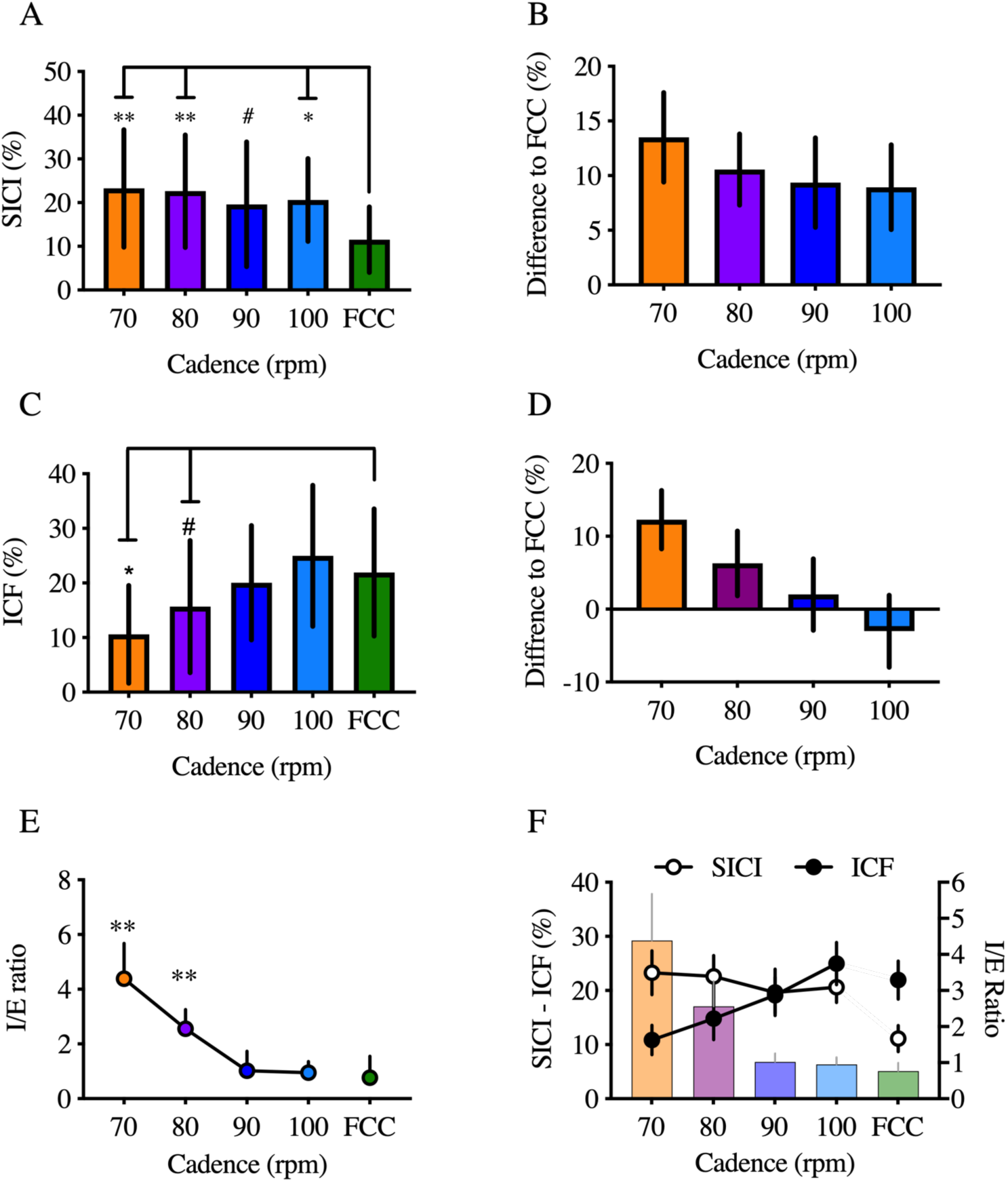
Group mean intracortical responses. **A:** Amount of SICI at each of the cadences. **B:** Percent difference in SICI at each of the fixed cadences compared to FCC. **C:** Amount of ICF at each of the cadences. **D:** Percent difference in ICF at each of the fixed cadences compared to FCC. **E:** I/E ratio at each of the cadences. **F:** SICI and ICF responses (open and filled circles respectively) together with the I/E ratio (right y-axes). **p < 0.01, *p < 0.05, #p = 0.07 depicts significant differences compared to the FCC.

### ICF

There was significant facilitation of the conditioned MEP compared to unconditioned MEP at all cadences (Table 1). There was a significant main effect of cadence on ICF (χ^2.^ = 14.03, df = 4, *P* = 0.007, Fig. 4C). ICF was less at 70 rpm compared to FCC (*P* = 0.012, r = 0.65), and there was no difference in ICF at 80, 90 and 100 rpm compared to FCC (*P* > 0.05).

### Inhibition-Excitation Ratio

There was significant main effect of cadence on I/E ratio (χ^2.^ = 12.44, df = 4, *P* = 0.014, Figure 4E). I/E ratio was higher at 70 (*P = 0*.*008, r = 0*.*65*) and 80 rpm (*P = 0*.*03, r = 0*.*49*) compared to FCC, with no difference at 90 (*P = 0*.*42*) and 100 rpm (*P = 0*.*32*) compared to FCC.

### Corticospinal Excitability (MEP)

Unconditioned MEP did not modulate with cadence (χ^2.^ = 3.53, df = 4, p = 0.48, Figure 4A) showing comparable levels of corticospinal excitability (70 rpm: 3.26 ± 0.41 mV, 80 rpm: 3.30 ± 0.31 mV, 90rpm: 3.32 ± 0.44 mV, 100 rpm: 3.01 ± 0.40 mV, FCC: 2.91 ± 0.97 mV). The coefficient of variation for all TMS pulses was 0.30 ± 0.04 and thus moderate.

### Maximal cycling EMG

There was no main effect of cadence on maximal cycling VL EMG (F_4,54_ = 0.89, p = 0.43) as EMG was comparable between cadences (Figure 5B).

**Figure 5.**
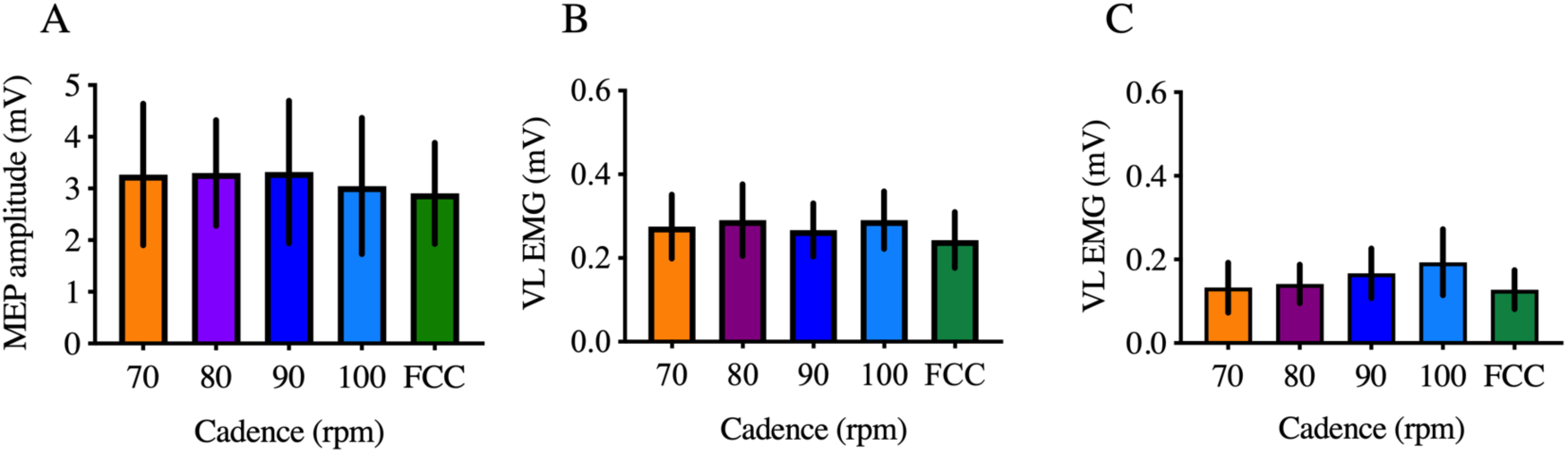
Corticospinal excitability and EMG responses. Panel **A** shows the mean peak to peak amplitude of the unconditioned MEP which was not different between the cadence. Panel **B** displays maximal VL EMG activation during cycling which was also not different between any of fixed cadences compared to the FCC. Panel **C** depicts the background EMG (bEMG) measured prior to each TMS stimulation which was not different between the cadences.

### Background EMG (bEMG)

There was no main effect of cadence on VL EMG prior to the stimulation (χ^2.^ = 2.48, df = 4, *P = 0*.*58*), demonstrating comparable levels of muscle activation at the time of stimulation (Figure 5C).

### Cadence

There was no difference between the fixed cadence of 70 and FCC (71.6 ± 8.1 rpm; *P* = 0.09).

## Discussion

### Main Findings

The main aim of this study was to investigate the effects of FCC on intracortical inhibitory and facilitatory circuits during constant submaximal workload leg cycling. The primary outcomes of the study demonstrate that SICI was reduced during FCC compared to a similar fixed cadence of 70 rpm; and ICF was augmented during FCC compared to a similar fixed cadence of 70 rpm. This suggests that intracortical mechanisms may contribute to the choice of cadence during locomotor cycling in humans, even though the chosen cadence is not one that is energetically optimal.

### Role of the motor cortex in cycling

In animals, rhythmical locomotor activities such as walking or running can be accomplished without the influence from supraspinal inputs. This is because an assembly of cells located in the spinal cord called central pattern generators (CPGs) generate rhythmical motor outputs (Grillner, 1981; Jordan, 1998). Similar to quadrupeds, it is believed that in humans, rhythmical motor outputs are also, at least partly, generated by spinal CPGs (e.g. Capaday *et al*., 1999; Pyndt & Nielsen, 2003; Carroll *et al*., 2006; Zehr *et al*., 2009), but that supraspinal input onto the spinal cord is crucial for movement control (Petersen *et al*., 2010; Sidhu *et al*., 2012; Forman *et al*., 2014). By using positron emission tomography (PET) and magnetic resonance imagining (MRI), studies have demonstrated that the motor areas of the cerebral cortex are active during walking (Fukuyama *et al*., 1997) and cycling (Christensen *et al*., 2000). Furthermore, non-invasive brain stimulation techniques such as subthreshold TMS have been used to demonstrate the contribution from intracortical inhibitory neurons in M1 (Davey *et al*., 1994; Butler *et al*., 2007) to the generation of locomotor drive during leg cycling (Sidhu *et al*., 2013). Our study provides supporting (i.e. SICI) and new (ICF) evidence for the role of the human motor cortex in cycling.

### Cycling cadence dependent effect on intracortical circuity

While there is evidence to show that cadence during human locomotor activity can influence sensory feedback – demonstrated by a suppression of somatosensory evoked potentials and diminished H-reflexes with increases in cycling cadence (Staines *et al*., 1997) – little is known of how cycling cadence influences the excitability of intracortical inhibitory and excitatory circuits. By using imaging techniques, it has been shown that there is a bilateral increase in the activity of the motor cortex with increasing pedaling frequencies and that this increase in activity is correlated with an increase in pedaling rate (Christensen *et al*., 2000). Using arm cycling, a recent study showed that when cadence increased, excitability of corticospinal neurons also increased (Forman *et al*., 2015). However, whether this modulation in M1 activation with cycling cadence is caused by changes in intracortical inhibitory and/or facilitatory mechanisms remains unclear. Our study provides new evidence to suggest that the increase in cortical activity with an increase in cycling cadence observed in previous studies using fMRI may be attributable to a decrease in SICI (e.g. increased activity of inhibitory interneurons) and an increase in ICF (Fig. 4C). Furthermore, we show that the balance between SICI and ICF (I/E ratio) is shifted towards a greater influence of SICI at lower cadences while the ratio approaches one at higher cadences – suggesting a balanced contribution of intracortical inhibition and facilitation at higher cadences of 90 and 100 rpm. This may have contributed to an increase in excitability of excitatory cortical neurons – reflected by an increase in ICF with increasing pedaling frequency (Fig. 4C). As such, it may be speculated that the balance between inhibition and excitation is a contributing factor as to why professional cyclists have higher preferred cadences (> 90rpm) compared to non-professional cyclists (Carnevale & Gaesser, 1991).

### Effect of freely chosen cadence on intracortical circuity

Despite the fact that the group average FCC of ∼72 rpm was not statistically different from the fixed cadence of 70 rpm, our study provides novel evidence to show that the magnitude of SICI and ICF was significantly different between the two conditions. Specifically, SICI was attenuated, while ICF was augmented during FCC compared to the fixed cadence of 70 rpm (Fig. 3B). Considering that the cadence, maximal cycling EMG and background EMG were not different between FCC and 70 rpm (Fig. 4B), as also shown in arm cycling studies (Marsh & Martin, 1995; Marais *et al*., 2004), it is unlikely that differences in sensory feedback contributed to the differences in SICI and ICF. Rather, the outcomes suggest that when subjects are prescribed to cycle at a cadence (even when this movement cadence is similar to their preferred movement cadence) the excitability of the intracortical inhibitory interneurons is augmented and the excitability of the intracortical facilitatory circuits is dampened. One consideration is the fact that when subjects are requested to cycle at a prescribed cadence, they shift the focus of their attention from an internal (freely chosen cadence) towards an external focus of attention (prescribed cadence). Adopting an external focus of attention has been shown to result in increased levels of SICI compared to an internal focus of attention (Kuhn et al. 2017). This is likely not an issue in the present study since after the FCC was established in each subject, they were asked to keep to their FCC via external feedback during the trial. It is possible that an internal model that represents augmented brain excitability is created to allow for preferred cadences during locomotor cycling.

While most previous studies have used fixed arm cycling cadences to investigate neural adaptations to power output (e.g. Forman *et al*., 2015; Lockyer *et al*., 2018), a more recent study investigated influence of self-selected versus prescribed cadence on corticospinal excitability in the arm muscles (Lockyer *et al*., 2019). In this study, subjects were asked to cycle at FCC in one session, and in the other session they had to cycle at a prescribed cadence of the same absolute cadence as the FCC; and during cycling they received single pulse TMS. Similar to our findings, they found no significant difference in MEPs during FCC versus a similar prescribed cadence. These findings suggest that the neural control of cycling cadence is likely similar between the upper and lower limbs, despite neurophysiological and anatomical differences.

### Neural mechanisms underlying SICI and ICF

It is believed that the neural mechanism underlying the reduction of the conditioned MEP after paired pulse stimulation during the SICI paradigm is similar to the subthreshold TMS causing a suppression of the ongoing EMG activity during walking (Petersen *et al*., 2001b) and cycling (Sidhu *et al*., 2013) – mediated by modulations in the excitability of the GABA_A_-ergic circuits within the primary motor cortex. Therefore, it is proposed that the increase in GABA_A_ inhibitory activity during non-preferred cadences is caused by a selective increase in the excitability of the GABA_A_-ergic circuits originating from L1 neurons which project on to the apical dendrites of pyramidal tract neurons and contribute to a selective suppression of the late I waves (Di Lazzaro & Rothwell, 2014). The origin of ICF, compared to SICI, is less clear because there is typically no modulation in the I-waves of the descending corticospinal volley associated with the facilitation of the MEP (Di Lazzaro *et al*., 2006; Ziemann, 2013). The rather broad range of the ISI between the subthreshold and suprathreshold MEP of 7-20 ms (Kujirai *et al*., 1993; Steve *et al*., 2006) suggests that ICF is mediated by neural networks separate from those involved in SICI. In particular, pharmacological studies show that ICF is mediated by NMDAreceptors (Hwa & Avoli, 1992) as the administration of NMDA receptor antagonists results in a decrease in ICF.

### Methodological and technical considerations

It should be acknowledged that the cadence mediated changes in ICF may not entirely reflect changes in cortical excitability. Apparently, ICF can be influenced by extracortical spinal effects (Wiegel *et al*., 2018). The authors of this work have shown that subthreshold TMS causes a facilitation of the H-reflex measured in the hand muscle flexor carpi radialis. In any case, this study also showed that there was no clear facilitation of the H-reflex by subthreshold stimulation when measured in the leg muscle soleus. Therefore, it is unlikely, although it cannot be entirely ruled out, that extracortical contributions influenced ICF during different cycling cadences in the current study.

Both SICI and ICF can modulate depending on the MEP size resulting from the test stimulation (Sanger *et al*., 2001; Opie & Semmler, 2014), which can inevitably confound the comparison of SICI and ICF between cadences. However, we did not see cadence dependent modulations in the control MEP size, as recently documented in the arm muscles (Lockyer *et al*., 2019). Therefore, the influence of any changes in test MEP response on SICI and ICF can be excluded. On the flip side, although we observed an increase In ICF with increasing cadence, this did not influence the net corticospinal excitability (i.e. magnitude of MEP). It is possible that the two intracortical (facilitatory and inhibitory) mechanisms and corticospinal excitability are not necessarily dependent on each other. There is in fact evidence to show that whilst intracortical inhibition modulates, the MEP does not necessarily do so (Singh et al., 2014; Smith et al., 2014). Also, by fixing the magnitude of the MEP response during a constant EMG contraction, change in intracortical inhibition is observed with interventions such as fatiguing exercise (Sidhu et al., 2018). These findings suggest that MEP and intracortical mechanisms may be mediated via independent mechanisms. It should also be acknowledged that the MEP is influenced by spinal mechanisms (McNeil *et al*., 2009; McNeil *et al*., 2013; Sidhu *et al*., 2018) and it is possible that spinal modulation with increasing cadence (i.e. increased spinal inhibitory input) influenced the net MEP response.

As described earlier, the measurements began with FCC, while all other cadences were randomized. Ideally, we would have randomized all cadences, but pilot testing showed that FCC was strongly influenced by the carry-over effects from the previous cadence. We also only recorded EMG activity from the agonist VL given that the focus of the study was on the extension phase of cycling. However, influences from the antagonist muscle activity may not be fully excluded and forms an important extension of the current study.

Finally, the present study did not include a cadence that was lower than the FCC (e.g. 60rpm). As such, the present study is not able to deduce if differences in SICI and ICF compared to the FCC would be apparent at lower cadences and should be explored in future work.

## Conclusion and significance

In conclusion, we provide new evidence that cortical physiologies including SICI and ICF modulate with cadence at a constant submaximal power output. More specifically, the present study provides new evidence to demonstrate that SICI is attenuated and ICF is augmented during FCC compared to a comparable prescribed cycling cadence.

## Additional Information

The author(s) declare no competing interests.

## Author contributions

S.S. and B.L. designed the experiment, B.L. analyzed the data, prepared figures and the initial draft of the manuscript, S.S. and B.L. reviewed and revised the manuscript. S.S. and B.L. agreed on the final version of the manuscript.

